# HOIL-1L deficiency induces cell cycle alteration which causes immaturity of myocyte and fibrogenesis

**DOI:** 10.1101/2023.09.29.560255

**Authors:** Kentaro Akagi, Shiro Baba, Hiroaki Fujita, Yasuhiro Fuseya, Daisuke Yoshinaga, Hirohito Kubota, Eitaro Kume, Fumiaki Fukumura, Koichi Matsuda, Takayuki Tanaka, Takuya Hirata, Kazuhiro Iwai, Junko Takita

**Author notes:** **Corresponding Author:** Shiro Baba, Address: 54 Kawahara-cho, Shogoin, Sakyo-ku, Kyoto City 606-8507, Kyoto, Japan.

## Abstract

HOIL-1L deficiency was recently reported to be one of the causes of myopathy and dilated cardiomyopathy (DCM). However, the mechanisms by which myopathy and DCM develop have not been clearly elucidated. Here, we sought to elucidate these mechanisms using the murine myoblast cell line C2C12 and disease-specific human induced pluripotent stem cells (hiPSCs). Myotubes were differentiated from control and HOIL-1L-KO C2C12 cells. Cardiomyocytes (CMs) were differentiated from control and patient-derived hiPSCs. We investigated the impact of HOIL-1L on differentiation of myotubes and CMs. Myotubes differentiated from HOIL-1L-KO C2C12 cells exhibited deteriorated differentiation and mitotic cell accumulation. CMs differentiaed from patient-derived hiPSCs had an abnormal morphology with a larger size and were excessively multinucleated compared with CMs differentiaed from control hiPSCs. Further analysis of hiPSC-derived CMs showed that HOIL-1L deficiency caused cell cycle alteration and mitotic cell accumulation. These results were supported by RNA sequencing of C2C12 cell-derived myotubes. In addition, *SerpinE2*, a cardiac fibrogenesis gene, was significantly upregulated in CMs differentiaed from patient-derived hiPSCs. These results demonstrate that abnormal cell maturation and fibrosis possibly contribute to the development of DCM and myopathy. In conclusion, HOIL-1L is an important intrinsic regulator of cell cycle-related myotube and CM maturation and cell proliferation.

## Introduction

Dilated cardiomyopathy (DCM) is defined by the presence of a dilated ventricle with severe systolic dysfunction, which causes end-stage heart failure (1). The reported prognosis of pediatric DCM patients is extremely poor. Many cases require cardiac transplantation, and the reported 5-year transplantation-free survival rate is only 55–63% (2, 3). The main evidence-based pharmacological therapies for DCM are digoxin, diuretics, angiotensin-converting enzyme inhibitors, and beta-adrenergic receptor blockers. These drugs improve the prognosis of adult DCM patients (4, 5); however, it is unclear whether they improve the prognosis of younger patients. One reason for this is that a wide variety of causes and associations have been described for DCM (6). These pathogenic categories include primary and secondary DCM, and the prognosis depends on them (7). Therefore, it is important to elucidate the mechanisms underlying each category of DCM to choose appropriate medical therapies.

Regarding progressive severe DCMs, HOIL-1L deficiency was recently reported to be one of the causes of pediatric DCM, and these patients also have myopathy, autoinflammatory syndrome, and pyogenic bacterial diseases (8). Most HOIL-1L-deficient patients are diagnosed with severe cardiomyopathy until adolescence. Cardiomyopathy is so severe and life-threatening that most of these patients require a heart transplantation in childhood. Unfortunately, heart transplantations are not performed for all patients because of problems including a lack of donor hearts and economic issues. Thus, it is crucial to develop treatments that affect the mechanisms responsible for the development of cardiomyopathy. The mechanisms by which cardiomyopathy develops in HOIL-1L-deficient patients have not been revealed. However, amylopectinosis in skeletal muscle and cardiomyocytes (CMs) of these patients has been identified by autopsy (8–10).

HOIL-1L is a component of the linear ubiquitin chain assembly complex (LUBAC), together with HOIP and SHARPIN. The LUBAC specifically generates a linear ubiquitin chain, which is formed via amino-terminal Met-1-linked ubiquitination (11–13). The linear ubiquitin chain plays an important role in mediating NF-κB activation and protecting against cell death (12, 13). Among the three components of the LUBAC, HOIL-1L, especially its UBL domain located in the N-terminal region, is essential for stability of the LUBAC (14) and linear ubiquitination at the TNFR1 signaling complex (15). Furthermore, the RING-IBR-RING (RBR) domain of HOIL-1L modulates immune signaling and cell death via monoubiquitination of the LUBAC (16). These facts shed light on the various clinical phenotypes of HOIL-1L-deficient patients, who also have autoinflammatory syndrome and pyogenic bacterial diseases. However, the mechanisms by which lethal cardiomyopathy develops in these patients have not been elucidated.

Recently, several DCMs were successfully modeled by generating patient-specific human induced pluripotent stem cell (hiPSC)-derived CMs (17–19). However, use of hiPSCs to model and study DCM in HOIL-1L-deficient patients has not been reported. Many HOIL-1L-deficient patients are diagnosed with DCM until childhood, and some cases develop DCM in infancy. Almost all HOIL-1L deficient patients have myopathy. Taking these facts into consideration, investigation of human cardiac cell development using hiPSC-derived CMs may be useful to elucidate the mechanism by which DCM develops upon HOIL-1L deficiency, and myoblasts can model the pathophysiology of HOIL-1L deficiency. In this study, we examined the role of HOIL-1L using murine myoblasts and hiPSC-derived CMs to reveal the mechanisms underlying myopathy and cardiomyopathy in HOIL-1L-deficient patients.

## Results

### HOIL-1L deletion perturbs murine myotube differentiation

To investigate the functions of the mammalian *HOIL-1L* gene in skeletal muscle, we used the murine myoblast cell line C2C12. HOIL-1L-KO C2C12 cells were generated using the lenti-CRISPR system. Mutation sites that cause myopathy and severe DCM, which necessitate intensive care of patients, were chosen, and models with KO at two locations, Exon 5 and Exon 7, were generated (Table 1). Exon 5 KO does not conserve the N-terminal domain, while Exon 7 KO does (Fig. 1A, B). Amylopectinosis was evaluated by measuring the amount of glycogen in myotubes differentiated from C2C12 cells on day 5 of differentiation *in vitro*. The amount of glycogen in myotubes differentiated from control, Exon 5-KO, and Exon 7-KO C2C12 cells did not significantly differ. Next, the fusion index and myosin heavy chain (MHC) density, which are indicators of differentiation, were calculated. The fusion index and MHC density on day 5 of differentiation were significantly lower in myotubes differentiated from Exon 5- and Exon 7-KO C2C12 cells than in those differentiated from control C2C12 cells (Fig. 1C). The protein expression level of MHC was also significantly lower in myotubes differentiated from Exon 5- and Exon 7-KO C2C12 cells than in those differentiated from control C2C12 cells (Fig. 1D). To assess the impaired phase of myotube differentiation, quantitative PCR (qPCR) was performed to measure the expression levels of the key myogenic transcription factors *Myogenin*, *MyoD*, and *MRF4*. *Myogenin* and *MyoD* expression levels in myotubes differentiated from control, Exon 5-KO, and Exon 7-KO C2C12 cells did not significantly differ during differentiation. However, the expression level of *MRF4,* an early differentiation factor, on days 5 and 6 of differentiation was significantly lower in myotubes differentiated from Exon 5- and Exon 7-KO C2C12 cells than in those differentiated from control C2C12 cells (Fig. S1). Collectively, these results indicate that HOIL-1L is one of the key factors that affect the early phase of murine myotube maturation. Furthermore, RNA sequencing (RNA seq) was performed of myotubes differentiated from HOIL-1L-KO and control C2C12 cells on day 5 of differentiation (Fig. 2A). The expression levels of 396 genes significantly differed by 2-fold or more between myotubes differentiated from Exon 5-KO C2C12 cells and those differentiated from control C2C12 cells (224 upregulated and 172 downregulated). Among these, expression of genes associated with muscle contraction and muscle structure (*ATP1B2*, *MB*, *TNNC2*, etc.) was significantly higher in myotubes differentiated from control C2C12 cells than in those differentiated from Exon 5-KO C2C12 cells (Fig. 2B). Gene set enrichment analysis (GSEA) and Gene Ontology (GO) analysis showed that the expression of genes linked to processes related to muscle contraction and myogenesis was also decreased in myotubes differentiated from Exon 5-KO C2C12 cells (Fig. 2C–E). Although expression of fewer genes was significantly changed by 2-fold or more in myotubes differentiated from Exon 7-KO C2C12 cells relative to those differentiated from control C2C12 cells, GSEA and GO analysis yielded the same results as those acquired with myotubes differentiated from Exon 5-KO C2C12 cells. These results indicate that HOIL-1L affects myotube differentiation regardless of deficient domain.

**Figure 1.**
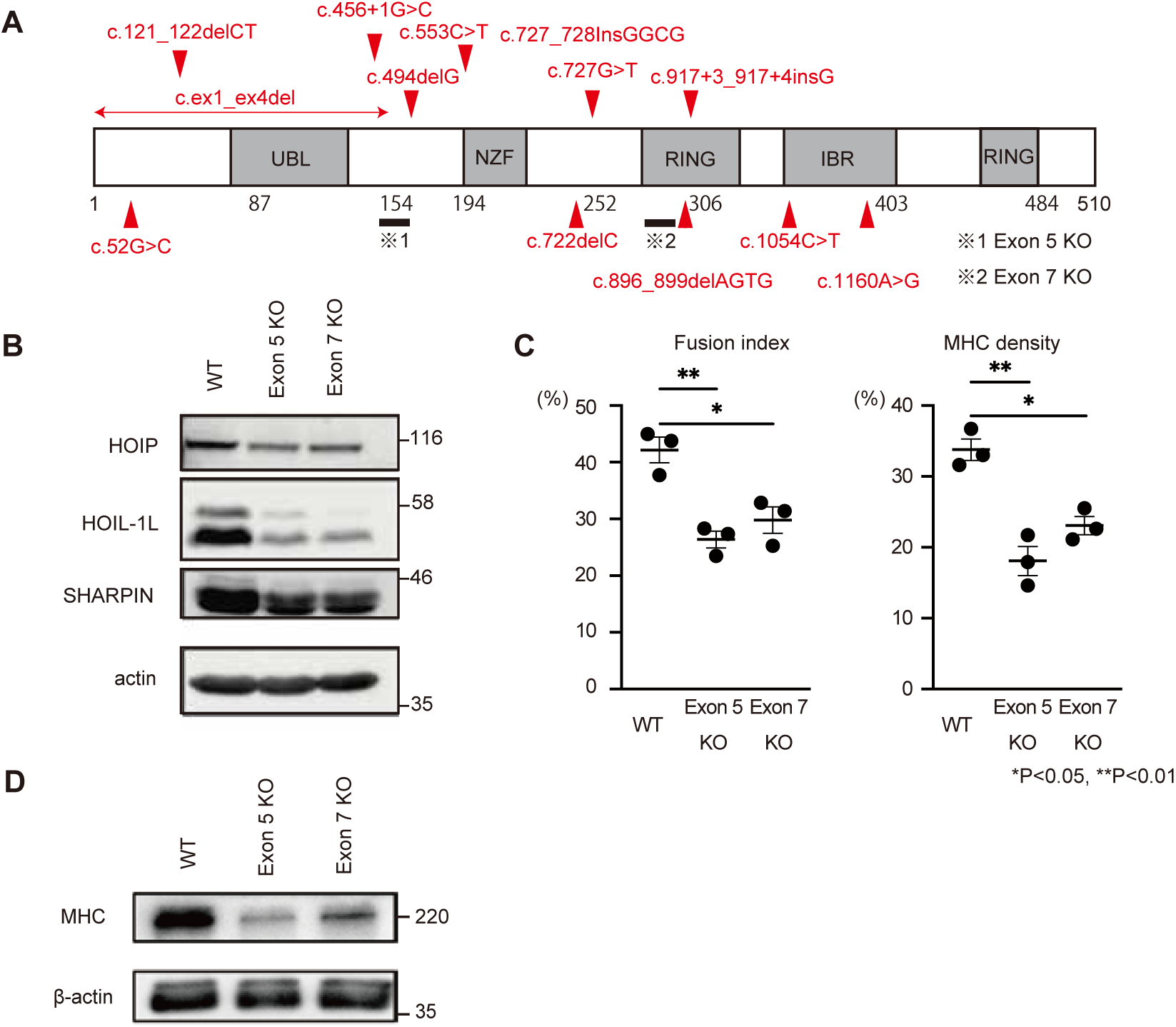
HOL-1L knockout C2C12 shows impaired myotube differentiation. **A**, Schematic presentation of reported mutations and HOIL-1L knockout locus of C2C12 myoblast in HOIL-1L protein domains. **B**, Immunoblot analysis of lysates of WT and HOIL-1L knockout myoblasts on day 5. Although HOIP and SHARPIN levels were decreased, HOIL-1L expression level was remarkably decreased in KO C2C12 derived myotubes. **C**, Fusion index and MHC density were calculated on day 5. All data are presented as mean±SEM. P values from Welch’s *t-test*. *P<0.05, **P<0.01 WT vs Exon 5 KO and Exon 7 KO. **D**, Western blotting shows impaired expression of MHC in HOIL-1L knockout C2C12 myotube.

**Table 1.** Type of mutations and main clinical phenotypes of reported HOIL-1L deficiency families.

| Reference | Patient | Allele1(DNA) | Allele2(DNA) | Allele1(Protein) | Allele2(Protein) | Exon | Clinical presentation |
| --- | --- | --- | --- | --- | --- | --- | --- |
| Boisson et al<br>2012<br>Nat Immunol | A | c.553C>T | c.ex1_ex4del | Q185* | 1-154 del(NTD) | E5/E1-4 | Immunodeficiency<br>Autoinflammation |
|  | B | c.121_122delCT | c.121_122delCT | L41fs*7 | L41fs*7 | E2 | **Cardiomyopathy** |
| Nilsson et al<br>2013<br>Ann Neurology | C | c.1160A>G | c.727G>T | N387S | E243* | E9/E6 | **Cardiomyopathy** |
|  | D | c.896-899delAGTG | c.896-899delAGTG | E299Vfs*18 | E299Vfs*18 | E7 | **Cardiomyopathy** → transplantation |
|  | E | c.722delC | c.722delC | A241Gfs*34 | A241Gfs*34 | E6 | **Cardiomyopathy** |
|  | F | c.52G>C | c.52G>C | A18P | A18P | E2 | **Cardiomyopathy** |
|  | G | c.ex1_ex4del | C.727_728InsGGCG | E243Gfs*114 | E243Gfs*114 | E6 | **Cardiomyopathy** → died of HF |
|  | H | c.1054C>T | c.1054C>T | R352* | R352* | E9 | **Cardiomyopathy** → transplantation |
|  | I | c.817+3_917+4 insG | c.817+3_917+4 insG | R298Rfs*40 | R298Rfs*40 | E7 | myopathy |
|  | J | c.494delG | c.494delG | R165Rfs*111 | R165Rfs*111 | E5 | **Cardiomyopathy** → transplantation |
| Krenn et al<br>2018<br>J Neurology | K and L | c.896-899delAGTG | c.896-899delAGTG | E299Vfs*18 | E299Vfs*18 | E7 | **Cardiomyopathy** → died of HF /<br>transplantation |

**Figure 2.**
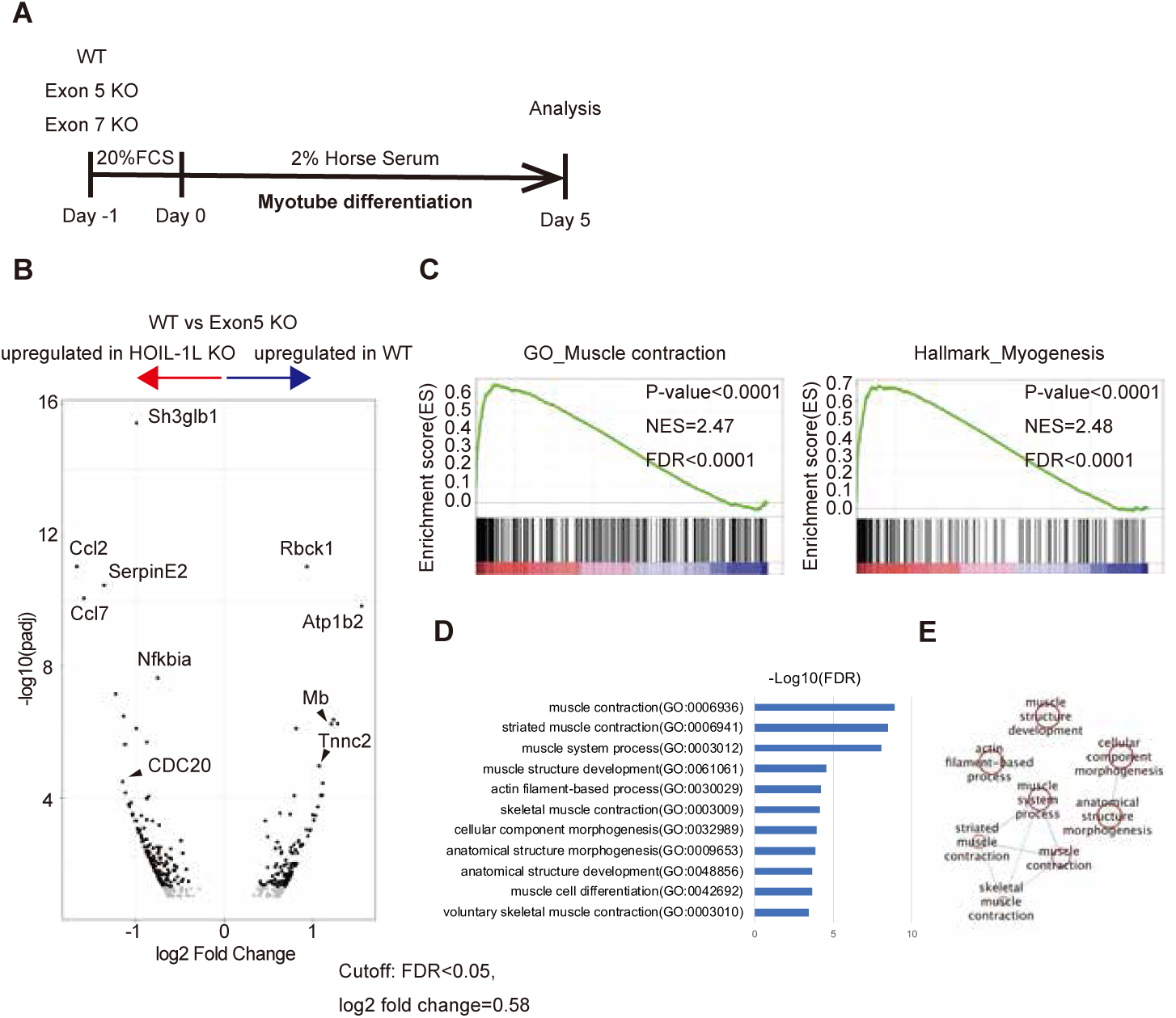
RNA-seq analysis of HOIL-1L KO C2C12 shows supportive data to implored myotube differentiation. **A**, A shame of myotube differentiation protocol of C2C12 myoblast. Myoblasts were cultured with DMEM containing 20% serum for 24h. On day 0, the medium was replaced to DMEM containing 2% horse serum and myoblasts were cultured for next 5 days for RNA-seq analyses. **B**, Distribution of log2 fold change between HOIl-1L Exon5 KO C2C12 myotubes and WT C2C12 myotubes versus the adjusted p-value (FDR). By setting the FDR cutoff as 0.1, 172 genes were upregulated in WT C2C12 myotubes and 224 genes were upregulated in HOIL-1L KO C2C12 myotubes. Genes associated with skeletal muscle function such as ATP1B2, MB and TNNC2 were down-regulated in HOIL-1L KO C2C12 myotubes. **C**, GSEA identified a significant enrichment of gene set associated with muscle contraction and myogenesis in WT C2C12 myotubes. **D**, the enrichment analysis by g: Profiler also showed significant enrichment of gene sets associated with muscle contraction and morphogenesis. **E**, Pathway enrichment analysis results are visualized in Cytoscape.

### HOIL-1L deletion alters the cell cycle according to RNA seq analysis of C2C12 myoblasts

Further GSEA revealed that genes associated with the G2M checkpoint and mitotic spindle formation were significantly upregulated in myotubes differentiated from HOIL-1L-KO C2C12 cells (Fig. 3A). Consistently, significantly more myoblasts were positively stained for phosphorylated (Ser10) histone H3, a marker of mitotic cells, in myotubes differentiated from HOIL-1L-KO C2C12 cells than in those differentiated from control C2C12 cells (Fig. 3B). *CDK1*, *AURKB*, *CDC20*, and *CCNB*2, which are involved in M phase of the cell cycle, were significantly upregulated in myotubes differentiated from HOIL-1L-KO C2C12 cells (Fig. 3C). These results suggest that cell cycle alterations induced by HOIL-1L deletion impair myotube differentiation.

**Figure 3.**
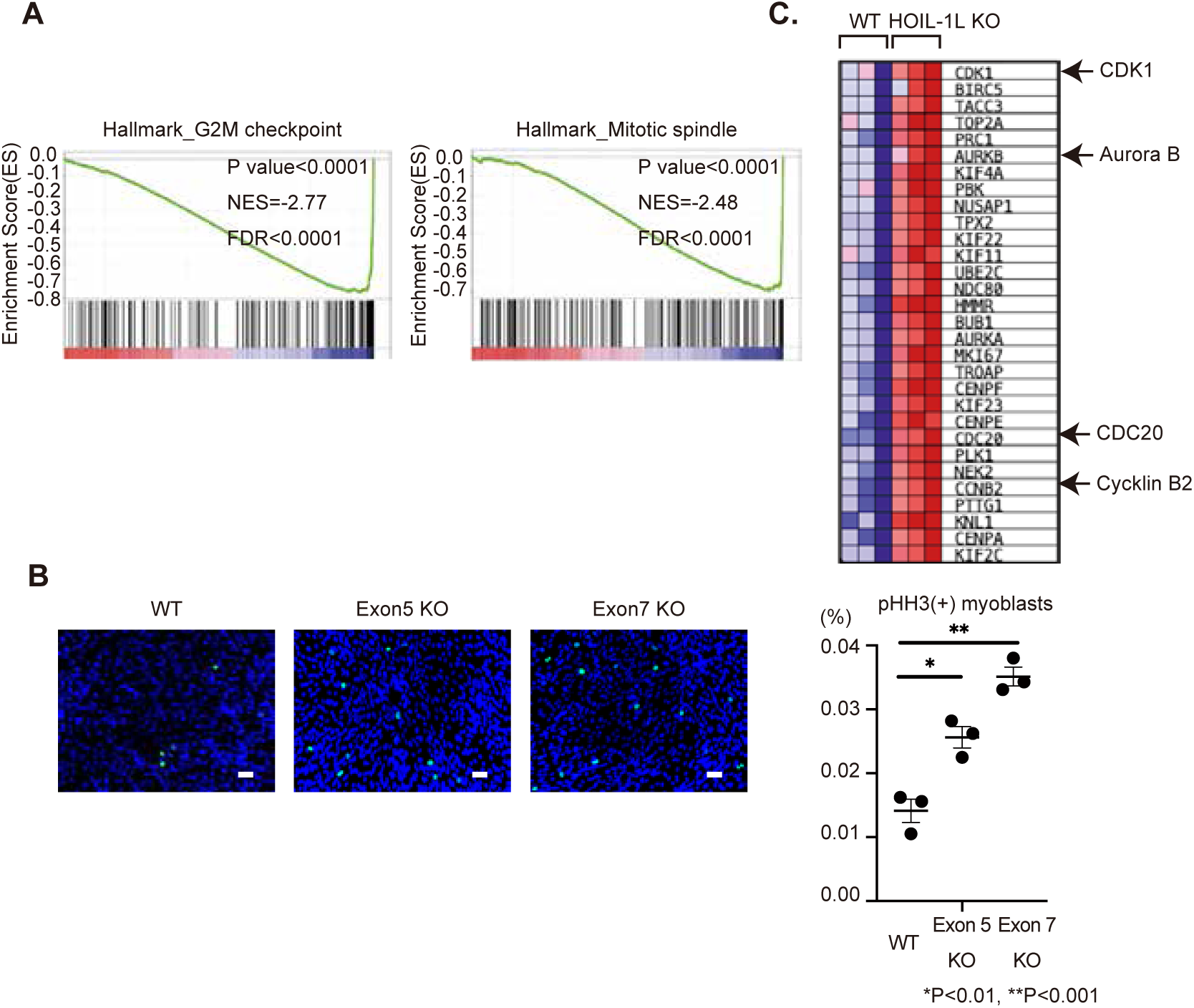
RNA-seq analysis of HOIL-1L KO C2C12 shows mitotic accumulation in HOIL-1L KO C2C12 myotubes. A, GSEA identified a significant enrichment of gene set associated with G2M checkpoint and mitotic spindle in HOIL-1L KO C2C12 myotubes. B, Immunostaining of histone H3 phosphorylation (pHH3) of C2C12 myoblasts showed more PHH3 positive HOIL-1L KO C2C12 myoblasts. Bar=50μm. All data are presented as mean±SEM. P values from Welch’s *t-test*. *P<0.05, **P<0.01 WT vs Exon 5 KO and Exon 7 KO. *P<0.01, **P<0.001. C, Heatmaps of the G2M check point associated genes enriched in HOIL-1L KO C2C12 myotubes compared with WT C2C12 myotubes. Especially, CDK1, AURKB, CDC20 and CCNB2, which promote entry into mitosis and maintain of the mitosis state.

### Clinical history of a HOIL-1L-deficient patient and hiPSC production

Next, we focused on the effect of HOIL-1L on CMs using hiPSCs. We generated hiPSCs from a HOIL-1L-deficient patient and two unrelated healthy control volunteers. Two hiPSC clones were established from the HOIL-1L- deficient patient. This patient was diagnosed with a homozygous missense mutation, a substitution of leucine for proline at residue 114 (L114P), by whole genome sequencing (Fig. 4A). She had an autoinflammatory syndrome, pyogenic bacterial diseases, and frequent ventricular arrhythmia with elevation of type-B natriuretic peptide (BNP), which possibly implied an initial presentation of heart failure. In infancy, she began to suffer from recurrent episodes of fever with lymphadenopathy. Based on laboratory examination, she was diagnosed with IgA deficiency at 2 years old and with IgG2 and IgG4 deficiency at 5 years old. During early childhood, she often suffered from bacterial pneumonia and otitis media, and from fever associated with lymphadenopathy and inflammatory bowel disease. In the second decade of life, she was diagnosed with epilepsy based on myoclonus seizures and spike-and-wave on an electroencephalogram, and she needed antiepileptic drugs to control the seizures. In the third decade of life, she developed a walking difficulty due to mild myopathy and palpitations due to ventricular arrhythmia in more than 15% of total heartbeats. Furthermore, echocardiography showed a mildly/moderately reduced left ventricular ejection fraction (51%) and a thin left ventricular wall with an elevated BNP level. She died in young adulthood because of these complications. Her elder sister, who possibly had the same mutation, was diagnosed with DCM and died of heart failure in young adulthood (Table S1). After performing pluripotency marker analysis by immunohistochemistry (Fig. 4B) and confirming that full-length HOIL-1L was not expressed in hiPSCs derived from the patient by western blotting (Fig. 4C), these hiPSCs were differentiated into CMs (hiPSC-CMs) and purified by glucose deprivation.

**Figure 4.**
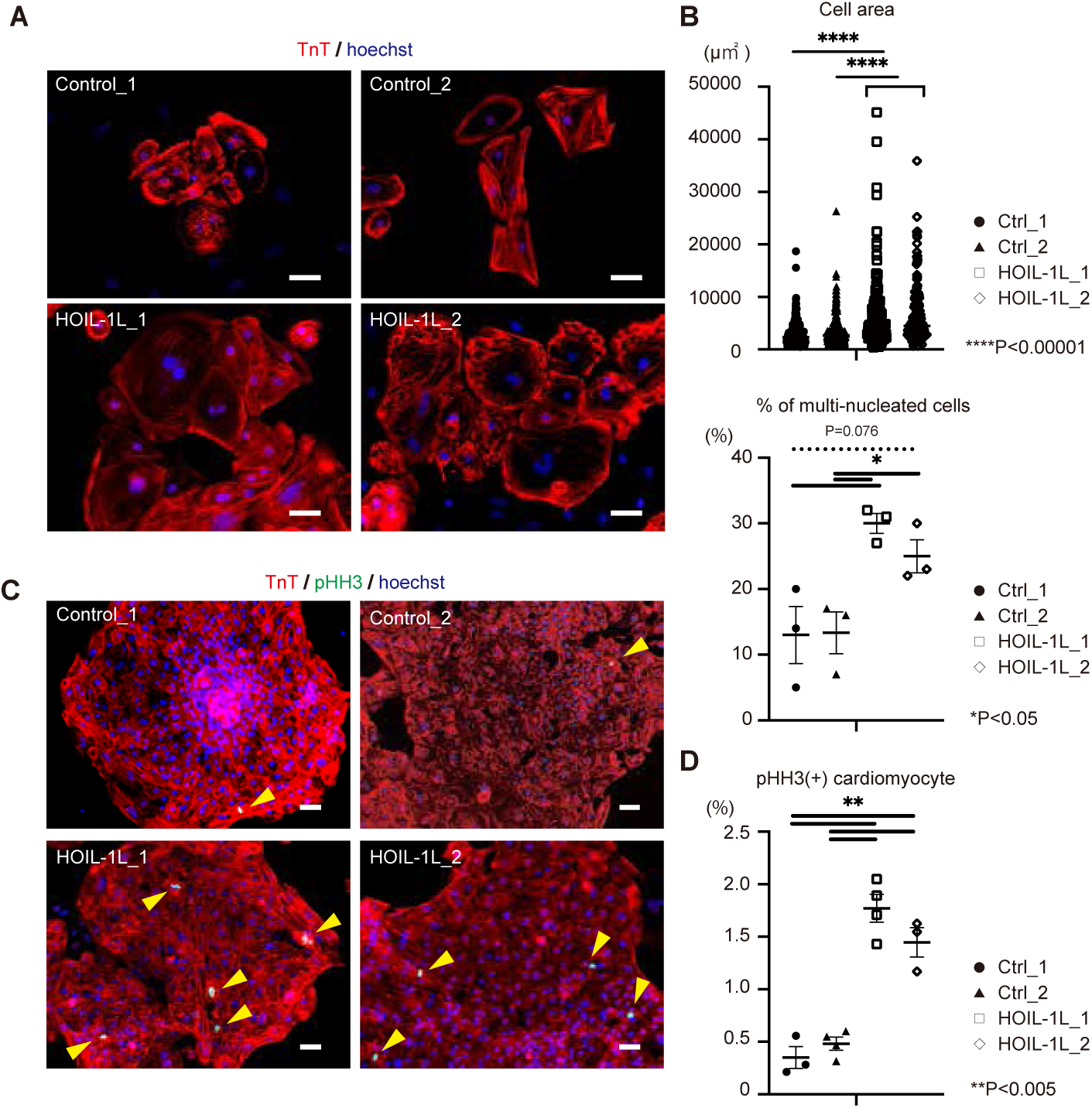
phenotype of HOIL-1L deficient patient-specific hiPSC-CMs. **A**, Representative immunofluorescence for cardiac troponin T (TnT). The patient hiPSC derived hiPSC derived cardiomyocytes showed larger size and multi-nucleation compared with control hiPSC derived cardiomyocyte. Bar=50μm. **B**, Quantification of cell size and multi-nucleation in control and HOIL-1L deficiency patient iPSC derived cardiomyocytes. (>different 3 passages and n>200, ****P<0.00001, *P<0.05) **C**, Immunostaining of histone H3 phosphorylation (pHH3) of hiPSC-CM showed more PHH3 positive hiPSC-CMs of HOIL-1L deficiency hiPSC-CMs than control hiPSC-CMs. Bar=50μm. **D**, Quantification of PHH3+ hiPSC-CMs showed significantly higher ratio in HOIL-1L deficiency hiPSC-CMs than control hiPSC-CMs. (*P<0.05, **P<0.005). All data are presented as mean±SEM. P values from Welch’s *t-test*.

### Amylopectinosis is not clearly detected in HOIL-1L-deficient hiPSC-CMs

To determine if the clinical features could be mimicked in patient-specific hiPSC-CMs (HOIL-1L hiPSC-CMs), amylopectinosis in the cytoplasm was evaluated by Periodic acid-Schiff (PAS) staining. All HOIL-1L and control hiPSC-CMs exhibited PAS-positive staining in their cytoplasm. However, this disappeared upon digestion with amylase in both HOIL-1L and control hiPSC-CMs. The glycogen level was not significantly higher in HOIL-1L hiPSC-CMs than in control hiPSC-CMs. (Fig. 4D, E). These results indicate that amylopectinosis cannot be reproduced in HOIL-1L hiPSC-CMs using a short period of *in vitro* differentiation and a culture protocol.

### HOIL-1L hiPSC-CMs are large and multinucleated

HOIL-1L hiPSC-CMs were significantly larger than control hiPSC-CMs. HOIL-1L hiPSC-CMs were more multinucleated than control hiPSC-CMs (Fig. 5A, B). RT-qPCR revealed that expression of *MYH7*, a cardiomyogenesis-related gene, was higher in HOIL-1L hiPSC-CMs than in control hiPSC-CMs (Fig. S2A). Based on these results, we hypothesized that HOIL-1L deficiency caused cytokinesis failure during cardiomyogenesis. In immunohistochemical analysis, significantly more HOIL-1L hiPSC-CMs than control hiPSCs-CMs were positively stained with phosphorylated (Ser10) histone H3, a marker of mitotic cells (Fig. 5C, D). Collectively, these results indicate that HOIL-1L deletion leads to abnormal cell cycle patterns in CMs and thereby causes excessive accumulation of these cells in M phase, resulting in production of larger CMs with more multinucleation.

**Figure 5.**
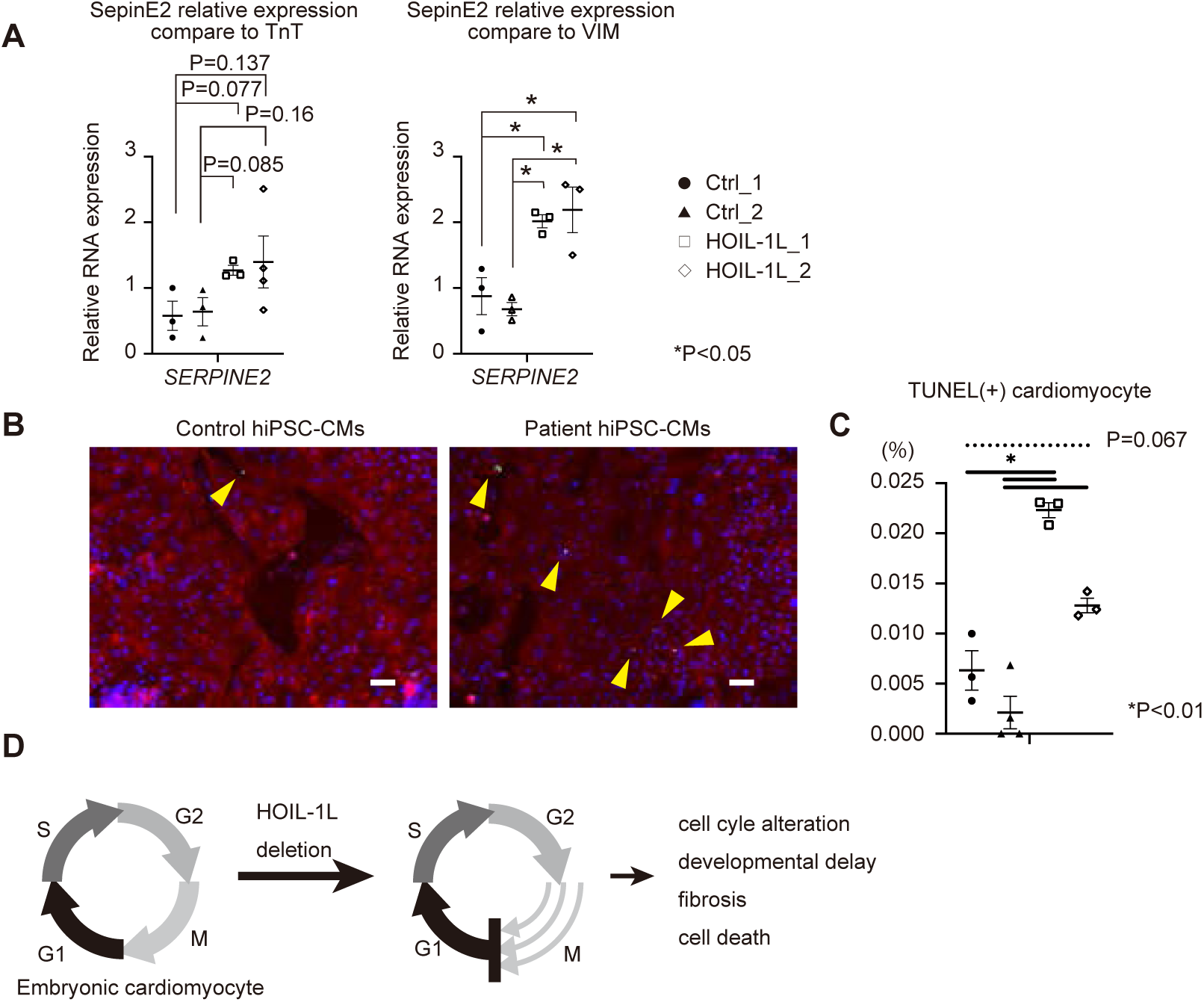
SerpinE2 expression in hiPSC-CMs and TUNEL assay. **A**, RT-qPCR of hiPSC-CMs show SerpinE2 overexpression in not cTnT positive HOIL-1L deficiency patient hiPSC-CMs, but VIM positive HOIL-1L deficiency patient hiPSC-cardiac fibroblasts. *P<0.05 **B**, Representative data of TUNEL staining of HOIL-1L deficiency hiPSC-CMs and control hiPSC-CMs. Bar=50μm. **C**, the ratio of TUNEL positive hiPSC-CMs was significantly higher in HOIL-1L deficiency compared to control. *P<0.01 **D**, Predictive mechanism of dilated cardiomyocyte development in HOIL-1L deficiency. All data are presented as mean±SEM. P values from Welch’s *t-test*.

### Heart development is perturbed in HOIL-1L-KO mouse embryos

To assess heart development *in vivo*, we investigated control and HOIL-1L-KO mouse embryos, which were generated previously (14). H&E staining of E10.5 normal, HOIL-1L null/+, and HOIL-1L null/null mouse embryos was performed. In normal embryos, the embryonic atrium and ventricle began to undergo septation to produce the four chambers. On the other hand, in HOIL-1L null/+ embryos, the heart began to form a round structure via ballooning. It was difficult to discriminate the atrium and ventricle during this period. In HOIL-1L null/null embryos, the heart structure was hardly detected (Fig. 6). These results indicate that HOIL-1L KO causes embryonic stunting including severe heart development delay.

### *SerpinE2* expression is elevated in HOIL-1L-KO cells

Lastly, we focused on the finding that expression of *SerpinE2* was significantly higher in myotubes differentiated from HOIL-1L-KO C2C12 cells based on RNA seq. *SerpinE2* is reportedly associated with cardiac fibrosis (29). In qPCR analysis, *SerpinE2* expression was significantly higher in HOIL-1L hiPSC-CMs than in control hiPSC-CMs when vimentin (*VIM*), a cardiac fibroblast gene, was used to standardize gene expression, but not cardiac troponin T (*cTnT*) (Fig. 7A). Expression of Col1a1, which leads to fibrosis, also tended to be higher in HOIL-1L hiPSC-CMs than in control hiPSC-CMs (Fig. S2B). These results indicate that fibrogenesis likely occurs in cardiac fibroblasts differentiated from HOIL-1L hiPSC-CMs compared with those differentiated from control hiPSC-CMs. To investigate fibrogenesis in detail, the effect of M phase accumulation and upregulated *SerpinE2* expression in hiPSC-CMs was evaluated by TUNEL staining. A significantly higher percentage of HOIL-1L hiPSC-CMs than control hiPSC-CMs was TUNEL-positive (Fig. 7B, C). These results indicate that accumulation in M phase causes death of CMs and that upregulated *SerpinE2* expression in cardiac fibroblasts causes paracrine fibrogenesis in differentiated cardiac cells (Fig. 7D).

## Discussion

DCM is a leading cause of heart failure, and the prognosis is poor. Heart transplantation is indicated for patients with end-stage DCM, but not all patients can undergo this procedure. While genetic alterations affect sarcomeric proteins and contractile function in some DCM cases, HOIL-1L deficiency is reportedly a novel cause of severe DCM. Mechanistic insight into this may help to develop a new treatment. For this purpose, development of hiPSC-CMs and the role of HOIL-1L in murine myoblasts were investigated in this study. Our results obtained using hiPSC-CMs and murine myoblasts indicate that cell cycle abnormalities probably contribute to the development of DCM in HOIL-1L- deficient patients.

In this study, we used patient-specific hiPSCs because recent advances in hiPSC technologies have made it possible to model genetic heart disease *in vitro*. Differentiated hiPSC-CMs could be purified to around 80–90% CMs, including atrial and ventricular cells, in experiments of cardiac differentiation. Therefore, a homogeneous population, the murine myoblast C2C12 cell line, was also used as a model of HOIL-1L myopathy, and this model showed the same phenotypes as HOIL-1L hiPSC-CMs. We performed experiments using both hiPSC-CMs and myotubes differentiated from C2C12 cells to focus not only on myopathy but also on cardiomyopathy. Lastly, all the data was gathered to analyze and discuss the disease mechanisms.

Amylopectinosis, an amylopectin-like polysaccharide, is observed in both skeletal and cardiac muscles in genetic diseases associated with glycogen metabolism (30). Amylopectinosis was detected by biopsy and autopsy of HOIL-1L-deficient patients (8–10, 31). However, we could not reproduce amylopectinosis in HOIL-1L hiPSC-CMs or myotubes differentiated from HOIL-1L-KO C2C12 cells. hiPSC-CMs reportedly exhibit metabolically immature phenotypes. These findings are expected because the main energy source is glycolytic metabolism in hiPSC-CMs but oxidative phosphorylation in matured CMs (32). Much longer culture and a more biogenic culture condition is presumably required to reproduce amylopectinosis.

C2C12 myoblasts spontaneously differentiate into myotubes upon differentiation in serum-free medium for 5–7 days, but myotubes do not exhibit spontaneous contraction during this differentiation. These facts are probably associated with our finding that amylopectinosis could not be reproduced. Thus, the effect of amylopectinosis in HOIL-1L-deficient patients remains unclear. If novel myocyte and CM differentiation protocols that mimic the long lifespans of mice and humans are developed, the effect of amylopectinosis on cardiomyopathy and myopathy will hopefully be revealed by *in vitro* experiments.

HOIL-1L hiPSC-CMs were morphologically abnormal, and differentiation of myotubes from HOIL-1L-KO C2C12 cells was impaired. These results imply that abnormalities in the maturation of CMs and skeletal muscle are related to the pathophysiology of DCM and myopathy in HOIL-1L-deficient patients. Hypertrophic vacuolar cardiomyopathy was reported to be a pathological finding in autopsies of HOIL-1L-deficient patients (31), and this is probably associated with impaired differentiation of CMs and development of DCM.

RNA seq revealed that expression of genes associated with muscle contraction and muscle structure was significantly lower in myotubes differentiated from HOIL-1L-KO C2C12 cells than in those differentiated from control C2C12 cells. This result supports our speculation.

Furthermore, GSEA detected increased expression of genes associated with mitosis regulation. *CCNB*2, which encodes a member of the cyclin family, was upregulated in myotubes differentiated from HOIL-1L-KO C2C12 cells. Members of the cyclin family play important roles in cell cycle control in CMs; therefore, HOIL-1L deletion was predicted to affect the cell cycle in CMs. To determine whether the cell cycle was abnormal in hiPSC-CMs, a mitotic cell marker was evaluated. This revealed that HOIL-1L hiPSC-CMs abnormally accumulated in M phase. During embryogenesis and fetal development of the mammalian heart, development of the cardiac chambers and further growth of the heart are accomplished by hyperplastic growth of CMs in the cell cycle. After birth, CMs exit the cell cycle and stop proliferating (33). Abrogation of the cell cycle results in multinucleation and polyploidy in CMs (34). Cell cycle alteration during late embryonic development due to loss of cytokinesis regulators (e.g., GAS2L3) leads to premature binucleation of CMs. GAS2L3 is required for mitosis of CMs around the late embryonic and early postnatal stages. Mice with loss of GAS2L3 exhibit compensatory CM hypertrophy, and this change is associated with interstitial fibrosis, ventricular dilation, and contractile dysfunction, resulting in early death of mice (35). Mitotic catastrophe, namely, mitotic cell accumulation, causes cell death (36). In patients with HOIL-1L deletion, CM hypertrophy, interstitial fibrosis, and ventricular dilation have been reported (8–10, 31), and these changes are thought to occur due to cell cycle alteration, accumulation of CMs in M phase, and eventually cell death. Recently, it was reported that LUBAC had a novel function in chromosome alignment and segregation. Interestingly, LUBAC mediated the promotion of CENP-E, kinetochore motor, attachment to kinetochores by liner ubiquitination (37, 38). Thus, the suppression of LUBAC causes a prolonged mitotic delay, chromosome missegregation during anaphase and increased mitotic cell death. The same result was revealed in our study, so further experiments are needed to show the relationship between cell cycle alteration in cardiomyocytes and the development of DCM in HOIL-1L deficiency.

In addition, SerpinE2 was significantly upregulated in myotubes differentiated from HOIL-1L-KO C2C12 cells. Subendocardial fibrosis has been reported in HOIL-1L-deficient patients (8, 31), and overexpression of SerpinE2 reportedly contributes to pathological cardiac fibrosis (29). SerpinE2 was significantly overexpressed in HOIL-1L hiPSC-CMs compared with control hiPSC-CMs. Based on these results, cardiac fibrogenesis is one of the mechanisms by which DCM develops in HOIL-1L-deficient patients.

The LUBAC is a trimer of HOIL-1L, SHARPIN, and HOIP, and has various functions such as in cell death and carcinogenesis. Cardiomyopathy and myopathy have been observed only in HOIL-1L-deficient patients; therefore, it is unlikely that SHARPIN, HOIP, and the LUBAC are also involved in these pathologies.

In conclusion, hypertrophic change of CMs observed at autopsy in HOIL-1L- deficient patients and developmental delay of the heart in HOIL-1L-KO mice were reproduced using HOIL-1L hiPSC-CMs, and cell cycle alteration was indicated to be one of the causes of these abnormalities. In addition, susceptibility to cardiac fibrosis also contributes to the onset and progression of DCM in HOIL-1L-deficient patients. Based on all these results, we conclude that HOIL-1L affects the development of myopathy and DCM. Our results will help to develop new treatments for lethal DCM in HOIL-1L-deficient patients. However, we could not explain the discrepancy between the phenotype of mice and human in this research. Further experiments are needed to reveal the difference between species.

## Methods

### hiPSC culture

hiPSCs were generated from an Asian HOIL-1L-deficient patient and healthy controls and kindly donated by the Center for iPS Cell Research and Application (Kyoto University, Kyoto, Japan). Patient-specific and healthy control hiPSCs were established using episomal vectors containing reprograming factors (20). Another control hiPSC line was established as described previously (21). Each cell line was maintained in mTeSR1 medium (Stem Technologies, Cat.# 85850) as previously reported (22). Cells were passaged every 4–5 days using accutase (Nacalai Tesque, Cat.# 12679-54). Dissociated cells were seeded on Matrigel-coated 6-well plates. The medium was supplemented with 5 μM Y27632 (TOCRIS, Cat.# 1254), a Rho-associated kinase inhibitor, on the first day of each passage.

### CM differentiation from hiPSCs

CMs were differentiated from hiPSCs using a previously reported protocol (23). Briefly, hiPSCs were seeded into a 12-well growth-factor-reduced (GFR) Matrigel-coated plate, grown for 4 days at 37°C in 5% CO_2_ and mTeSR1 medium, and allowed to reach 80–90% confluency. On day 0 of differentiation, the medium was changed to differentiation media, which was RPMI containing 2% B27 minus insulin supplement (Gibco, Cat.# A18956-01) and 10–12 μM CHIR99021 (Selleck, Cat.# S2924), a GSK3 inhibitor. After incubation for 24 hours, the medium was replaced with fresh differentiation medium. On day 3, the medium was replaced with differentiation medium containing 5 μM IWP-2 (TOCRIS, Cat.# 3533), a Wnt inhibitor. On day 5, the medium was replaced with fresh differentiation medium. On day 7, B27 minus insulin was replaced with a B27 supplement (Gibco, Cat.# 17504044). Differentiated hiPSC-CMs were purified in glucose-depleted lactate medium as described previously (24).

### C2C12 cell culture

C2C12 cells were kindly provided by Dr. Yuji Yamanashi (The Institute of Medical Science, The University of Tokyo) (25). The growth medium was DMEM/F12 (Sigma-Aldrich, Cat.# D6421) containing 20% FBS, 2 mM glutamine (Gibco, Cat.# 25030081), 100 units/mL penicillin, and 100 μg/mL streptomycin. Cells were incubated at 37°C in a humidified incubator containing 5% CO_2_. Myoblasts were differentiated into myotubes in DMEM/F12 medium containing 2% horse serum (Gibco, Cat.# 16050122, Lot. 1968945) (26, 27).

### Generation of HOIL-1L-KO C2C12 cells

Lenti-CRISPR v2 (Addgene, Cat. # 52961), which contains a puromycin resistance gene, carrying a guide RNA oligonucleotide (5’-acctcacccttcagtcacgg-3’ for Exon 5 of the *HOIL-1L* gene or 5’-acgcagcaccacggcctcgc-3’ for Exon 7 of the *HOIL-1L* gene) was constructed. HEK293T cells were transfected with the plasmids using Lipofectamine 2000 (Thermo Fisher, Cat.# 11668019). Viruses were harvested at 48 hours after transfection, and the media were filtered through a 0.45 μm PES filter. C2C12 cells were transduced with the viruses in medium containing 10 μg/mL polybrene. At 24 hours after transduction, puromycin selection was started. The selected cells were collected and KO of HOIL-1L was confirmed by Sanger DNA sequencing.

### Immunohistochemistry and TUNEL assay

Myotubes differentiated from C2C12 cells on day 5 of differentiation were fixed in 4% paraformaldehyde (PFA) for 1 hour at 4°C, permeabilized in 0.1% Triton X-100 for 10 minutes at room temperature, and blocked in PBS containing 3% skim milk for 1 hour. Thereafter, myotubes were stained with an anti-MHC antibody (1:200, mouse monoclonal, R&D Systems, Cat.# MAB4470). The fusion index was calculated by dividing the number of nuclei in myotubes by the total number of nuclei in a field of view (26). The MHC density was calculated by dividing the area occupied by MHC-positive myotubes by the total area of the field of view. The fusion index and MHC density were reported as averages of at least three fields of view (>500 total nuclei). Three independent experiments were performed for the calculation. For pluripotency marker analysis, undifferentiated hiPSC colonies were fixed in the same way, and fixed cells were stained with mouse anti-Oct3/4 (1:50, Santa Cruz Biotechnology, Cat.# <u>sc5279</u>) and anti-TRA1-81 (1:100, Millipore, Cat.# MAB4381) antibodies. Cells were then incubated with Alexa Fluor-conjugated secondary antibodies (1:1000) and Hoechst 33342. For immunofluorescence microscopy analysis of hiPSC-CMs, size and multinucleation were analyzed after around 50–60 days of differentiation and mitosis was analyzed after 20 days of differentiation. hiPSC-CMs were replated onto GFR Matrigel-coated 24-well dishes, incubated at 37°C in 5% CO_2_ for 72 hours, fixed in 4% PFA for 1 hour at 4°C, permeabilized in 0.1% Triton X-100 for 10 minutes at room temperature, and blocked in PBS containing 3% skim milk for 1 hour. Thereafter, hiPSC-CMs were stained with anti-cTnT (1:100, mouse monoclonal, Thermo Fisher, Cat.# MA5-12960) and anti-phospho-histone H3 (Ser10) (1:1000, rabbit monoclonal, Cell Signaling, Cat.# D7N8E) antibodies. After primary antibody treatment, cells were rinsed three times with PBS for 5 minutes at room temperature and then incubated overnight with secondary antibodies diluted 1:1000 in PBS. Nuclei were stained with Hoechst 33342 (1:1000, Invitrogen, Cat.# H3570). For isotype controls, mouse IgG1 isotype (BD Biosciences, Cat.# 554121) and rabbit IgG isotype (BD Biosciences, Cat.# 550875) were used. All immunofluorescence analyses were performed using a BZ-710X microscope (Keyence). The TUNEL assay was performed using a Cell Death Detection Kit (Roche, Cat.# 11684795910) following the manufacturer’s protocol.

### Western blotting

Myotubes at day 5 of differentiation were lysed in M-PER buffer (Thermo Scientific, Cat.# 78501) containing 1× protease inhibitor and then incubated on ice. The samples were sonicated on ice for 30 seconds. The lysates were incubated on ice for 10 minutes and then centrifuged at 15,000 rpm for 15 minutes. Protein concentrations were determined using the Bradford assay. Thereafter, 30 μg of protein was loaded onto each lane of 10% SDS-PAGE gels. The membranes were probed with an anti-MHC antibody (1:100, R&D Systems, Cat.# MAB4470) in blocking buffer (5% BSA) at 4°C overnight, washed, incubated in secondary antibodies for 1 hour at room temperature, developed using ECL western blotting substrate (Bio-Rad, Cat.# 1705060), and imaged using the ChemiDoc MP Imaging System (Bio-Rad).

### Flow cytometry

hiPSC-CMs were dissociated on the day of evaluation by incubating them in 0.25% trypsin-EDTA for 10–15 minutes at 37°C. They were fixed in Cytofix/Cytoperm solution (BD Biosciences, Cat.# 554714) for 20 minutes at 4°C, washed with BD Perm/Wash buffer (Cat.# 554723), stained with an anti-cTnT antibody (1:200, mouse monoclonal, Thermo Fisher, Cat.# MA5-12960) followed by Alexa Fluor-conjugated secondary antibodies, and analyzed using FACSverse (BD Biosciences). Data were collected from at least 10000 events. Data with >80% cTnT populations were used for all experimental analyses.

### qPCR analysis of C2C12 cell-derived myotubes and hiPSC-CMs

Total RNA was extracted from day 0 to day 7 of myotube differentiation using an RNeasy Mini Kit (Qiagen, Cat.# 74104) according to the manufacturer’s instructions. qPCR was performed using SYBR Green PCR Master Mix (Takara, Cat.# RR820) on a StepOnePlus system (Thermo Fisher Scientific) with the ΔΔCt method. GAPDH was used to standardize gene expression. Total RNA was extracted from hiPSC-CMs at 45–60 days after differentiation.

### H&E staining

HOIL-1L-KO and control mouse embryos were generated as described previously (14). Paraffin sections of E10.5 HOIL-1L null/+, HOIL-1L null/null, and control littermate mouse embryos were deparaffinized and stained with H&E.

### RNA seq

RNA was isolated from C2C12 cells using a RNeasy Mini Kit, according to the manufacturer’s instructions. RNA integrity was measured using an Agilent 2200 TapeStation and RNA Screen Tapes (Agilent Technologies). Sequencing libraries were prepared using a NEBNext Ultra II RNA Library Kit for Illumina (New England Biolabs) with the NEBNext Poly (A) mRNA Magnetic Isolation Module (New England Biolabs), according to the manufacturer’s protocol. Prepared libraries were run on an Illumina HiSeq X sequencing platform in 150 bp paired-end mode. Sequencing reads were aligned to the GRCm38 mouse genome assembly using STAR (c.2.5.3). Mapped reads were counted for each gene using the GenomonExression pipeline (https://github.com/Genomon-Project/GenomonExpression). Normalization of the read counts of RNA seq data and differential expression analysis were performed using the Bioconductor package DESeq2 (version 1.26.0). Differentially expressed genes with a greater than 2-fold change and a false discovery rate less than 0.1 were filtered and evaluated.

### GSEA

GSEA was performed using software (version 4.0.3) from the Broad Institute. Normalized expression data obtained from RNA seq were assessed using GSEA software and the Molecular Signature Database (http://www.broad.mit.edu/gsea/). c5 ontology gene sets were used, and a false discovery rate less than 0.01 was considered to be statistically significant. Pathway enrichment analysis using g:Profiler and visualization of enrichment results in an enrichment map were performed using Cytoscape software (version 3.7.2) as described previously (28).

### Statistics

Data are shown as mean ± SEMs, as indicated in the figure legends. All statistical analyses were performed using Welch’s *t-test* with GraphPad Prism (version 9.00, GraphPad Software). *p*<0.05 was defined as significant.

### Study approval

All experiments were conducted in accordance with the guidelines of the Institutional Ethical Committee of Kyoto University and the Declaration of Helsinki.

## Author Contributions

KA, SB, KI, and JT designed the experiments and the overall study. KA and SB wrote the manuscript. KA, HF, and YF generated HOIL-1L-KO C2C12 cells. KA, DY, KM, TT, and TH performed the experiments. KA and HK performed the bioinformatics analysis. All authors discussed the results, and helped to prepare and edit the article.

## Acknowledgments

We are grateful to Dr. Yuji Yamanashi for providing C2C12 cells, Kumi Kodama for assisting with preparation and performance of the experiments, and Dr. Tomohiro Morio for recruiting a patient. Preparation of paraffin-embedded sections and H&E staining were supported by the Anatomic Pathology Center of the Graduate School of Medicine of Kyoto University.

## Non-standard Abbreviations and Acronyms

LUBAC: linear ubiquitin chain assembly complex
hiPSC: human-induced pluripotent stem cell
hiPSC-CM: human-induced pluripotent stem cell-derived cardiomyocyte

## Novelty and significance

What is known?

- HOIL-1L deficiency has been reported to be one of the causes of pediatric dilated cardiomyopathy (DCM). Although it’s a lethal phenotype, the mechanism of developing DCM in HOIL-1L deficiency has not been revealed at all.
- Some pathological findings, amylopectinosis and interstitial fibrosis in cardiomyocyte, have been reported, and they can be clues for elucidation of the mechanism.

What new information does this article contribute?

- We found the impaired cardiomyocyte maturation and alteration of cell cycle in HOIL-1L patient hiPSCs-derived cardiomyocytes, providing a clue of the mechanism of developing DCM in HOIL-1L deficiency.
- Furthermore, SerpinE2 has been recognized as one of the reason of cardiac fibrosis. This fact will lead to the development of treatments for DCM in HOIL-1L deficiency patients.

---

**Fig. S1.**
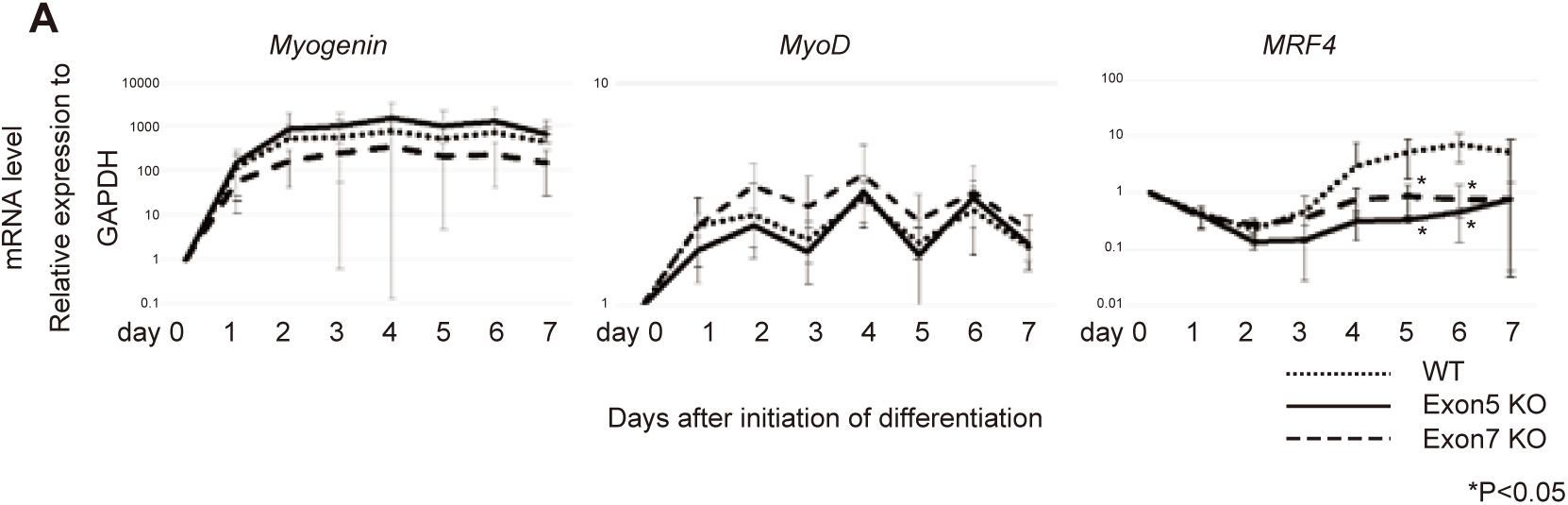
Gene expression during myotube differentiation in C2C12. **A,** MRF4 expressions were significantly decreased in HOIL-1L knockout C2C12 myotubes. All data are presented as mean±SEM. P values from Welch’s *t-test*. (*P<0.05 WT vs Exon 5 and Exon 7 KO.)

**Fig. S2.**
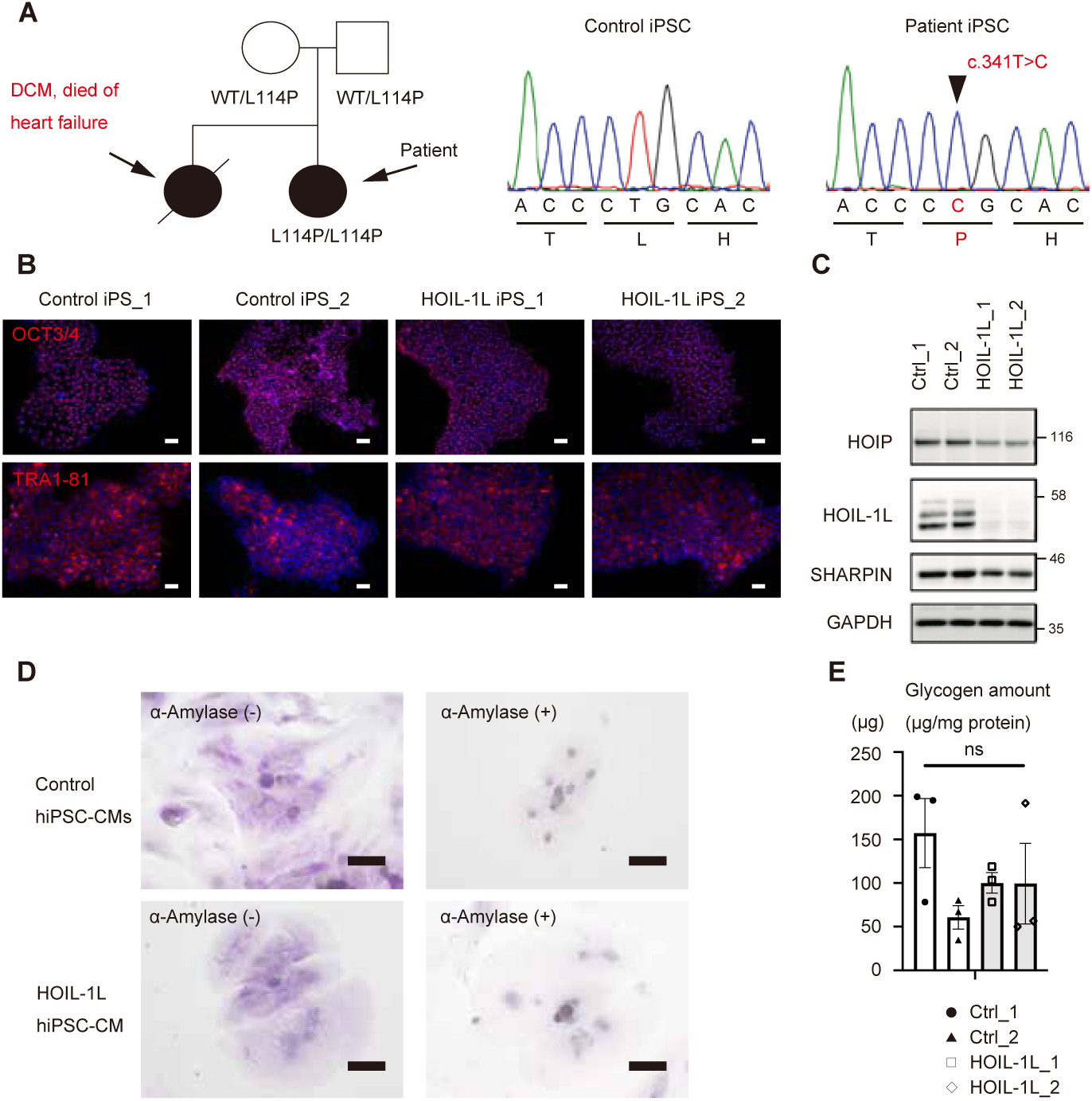
Characterization of HOIL-1L deficient patient-specific hiPSC-CMs and amylopectinosis assay in hiPSC-CMs. **A,** Schematic pedigrees of a family with HOIL-1L deficiency. The patient is indicated by arrow, and her old sister died of cardiac failure. Her parents have heterozygous L114P missense mutation on exon 4 of the HOIL-1L gene and the patients has homozygous L114P missense mutation on HOIL-1L gene. Two cell lines were established, and L114P (c.341T>C) were confirmed by sequence analysis of PCR-amplified genomic DNA. **B,** Immunofluorescence for OCT3/4 and TRA-1-81 of representative hiPSC colonies of controls and the patient. **C,** Western blotting analysis of lysates of hiPSC from control, patient specific and HOIL-1L KO. **D,** Amylopectinosis, which is not able to be digested by α-amlyase, was not detected both in control and HOIL-1L deficiency hiPSC-CMs. Bar=50μm **E,** Glycogen amounts of Control and HOIL-1L deficiency hiPSC-CMs were not significantly different, either.

**Fig. S3.**
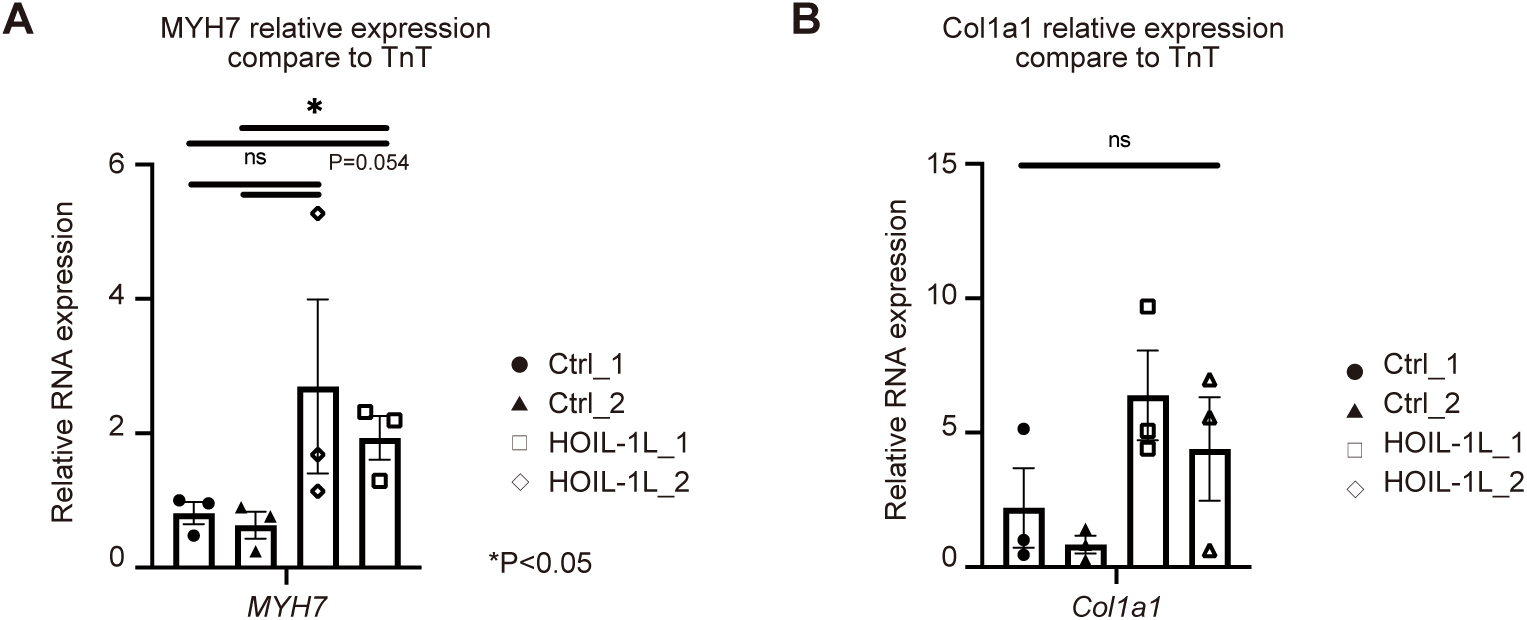
Analysis of MYH7 and Col1a1 by RT-qPCR. **A,** RT-qPCR of hiPSC-CMs show higher MYH expression in HOIL-1L deficiency patient hiPSC-CMs than control hiPSCs-CMs. **B,** the expression of Col1a1, which leads fibrosis, tend to higher in HOIL-1L deficiency patient hiPSC-CMs than control hiPSCs-CMs. All data are presented as mean±SEM. P values from Welch’s *t-test*.

**Figure S4.**
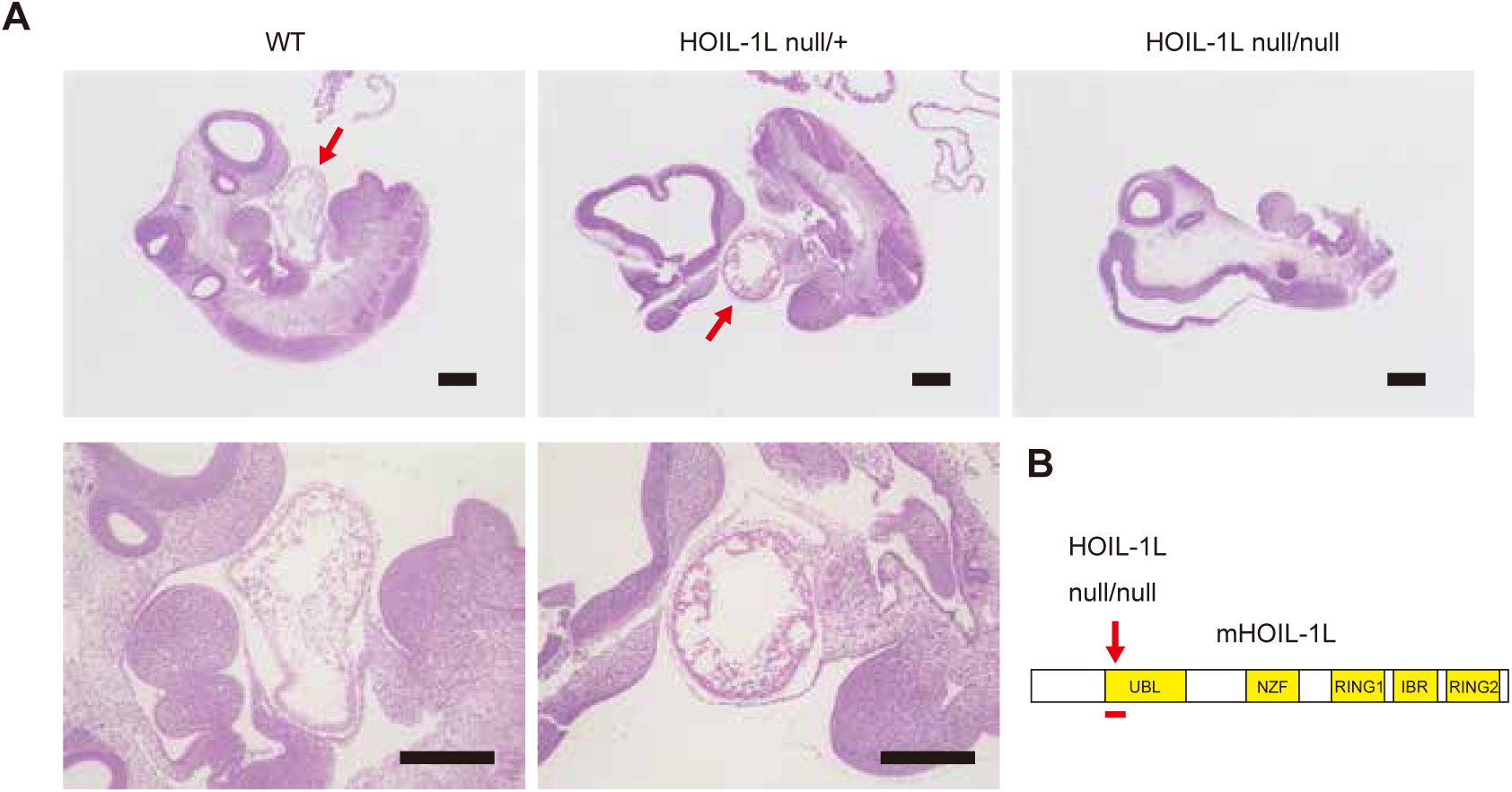
Hematoxylin-eosin staining of HOIL-1L KO fetal mouse. **A,** Representative image of Hematoxylin-eosin staining on E10.5 whole embryo paraffin embedded sections. A wild type mouse embryo heat developed normally (red arrow of left upper panel, magnified image is presented in left lower panel). The section of HOIL-1L null/+ hetero mouse shows obviously delayed cardiac development (red arrow in middle upper panel, magnified image is presented in middle lower panel). HOIL-1L null/null mouse shows completely defect of cardiac development accompanies with embryonic lethality (right upper panel). Bar=300μm **B,** Schematic representation of HOIL-1L null/null mutation from Fujita et al (2018). *Cell Report 23, 1192-1204*

**Table S1.** Clinical features of the HOIL-1L deficiency patient.

| Feature | Description |
| --- | --- |
| Growth | Normal |
| Immunological | High frequency of cervical lymphadenitis and otitis media<br>→diagnosed as IgA deficiency (2 years old), IgG2 and IgG4 deficiency (5 years old) |
| Gastrointestinal | Recurrent diarrhea and gastroenteritis |
| Cardiac | Frequent premature ventricular contractions (20,000-40,000 beats/day) and elevated BNP (68.9pg/ml) |
| Musculoskeletal | Walking difficulty by mild myopathy |
| Neurologic | Epilepsy (myoclonus seizures controled by antiepileptic drugs) |
| Familial history | Her old sister were diagnosed as dilated cardiomyopathy, and died of heart failure at 37 years old. |

